# Multiuse patient monitoring equipment as a risk factor for acquisition of *Candida auris*

**DOI:** 10.1101/149054

**Authors:** Hilary Madder, Ian Moir, Ruth Moroney, Lisa Butcher, Robert Newnham, Mary Sunderland, Tiphanie Clarke, Dona Foster, Peter Hoffman, Ginny Moore, Colin S Brown, Katie JM Jeffery

**Author notes:** Joint first authors (Hilary Madder, Ian Moir, Ruth Moroney).

## Introduction

*Candida auris* is an emerging and multi-drug resistant pathogen recently associated with outbreaks worldwide, often in intensive care settings (1). Following the alerts issued by the CDC and PHE in June 2016 (2,3), a look back exercise in the neurological intensive care unit (NITU) in the Oxford University Hospitals NHS Foundation Trust identified five clinical cases of *C. auris* infection. After the identification of an additional isolate in a urine sample, a patient and environmental screening programme was introduced in the last week of October 2016. We report here a summary of our initial findings which other affected units may wish to consider as part of their infection prevention and control measures.

## Methods

### Patient screening

Patients were screened on admission to the NITU, weekly, and on discharge, from October 2016 to May 2017. Screening sites were nose, axilla, groin, tracheostomy and wound swabs (Sigma Transwab®), and urine culture. This was modified in January 2017 to three times weekly screening via axilla and groin swabs, with a full specimen screen on admission and discharge. Patients on the neuroscience ward (the main step-down ward for NITU) were also screened weekly. Swabs were cultured on Sabouraud Dextrose Agar W/Chloramphenicol (SabC) at 37°C in air. Colonies with morphological characteristics of *C. auris* were identified using via the Bruker MALDI biotyper system - resulting spectra were analysed with MALDI biotyper RealTime Classification software version 3.0.

### Environmental screening

Various methodologies for environmental screening were used with an emphasis on sampling high-touch areas and multi-use devices. These included the use of bacterial swabs in a liquid transport medium (Sigma Transwab®), sponges to sample larger surface areas (Polywipe^™^), and Sabouraud dextrose agar contact plates (55mm diameter). Enrichment culture was performed in brain heart infusion or malt extract broth. The air was sampled using passive air sampling techniques (SabC settle plates) and two active air samplers (an AirIdeal and Coriolis air sampler operating at 100 and 300 L/min respectively). Confirmation of *C. auris* was achieved using the Bruker MALDI biotyper system as above.

## Results

### Patient screening

Between the end of October 2016 to the end of May 2017, approximately 15,000 screens from 726 individual patients were received. Two hundred and eighty patients were from NITU, 407 from the neuroscience ward and 39 from other clinical areas, including the adult intensive care unit. Fifty-eight patients became *C. auris* colonised or infected, with two cases of candidaemia (Figure 1). Fifty-seven of the 58 patients colonised or infected with *C. auris* had been admitted to NITU during their hospital stay.

**Figure 1.**
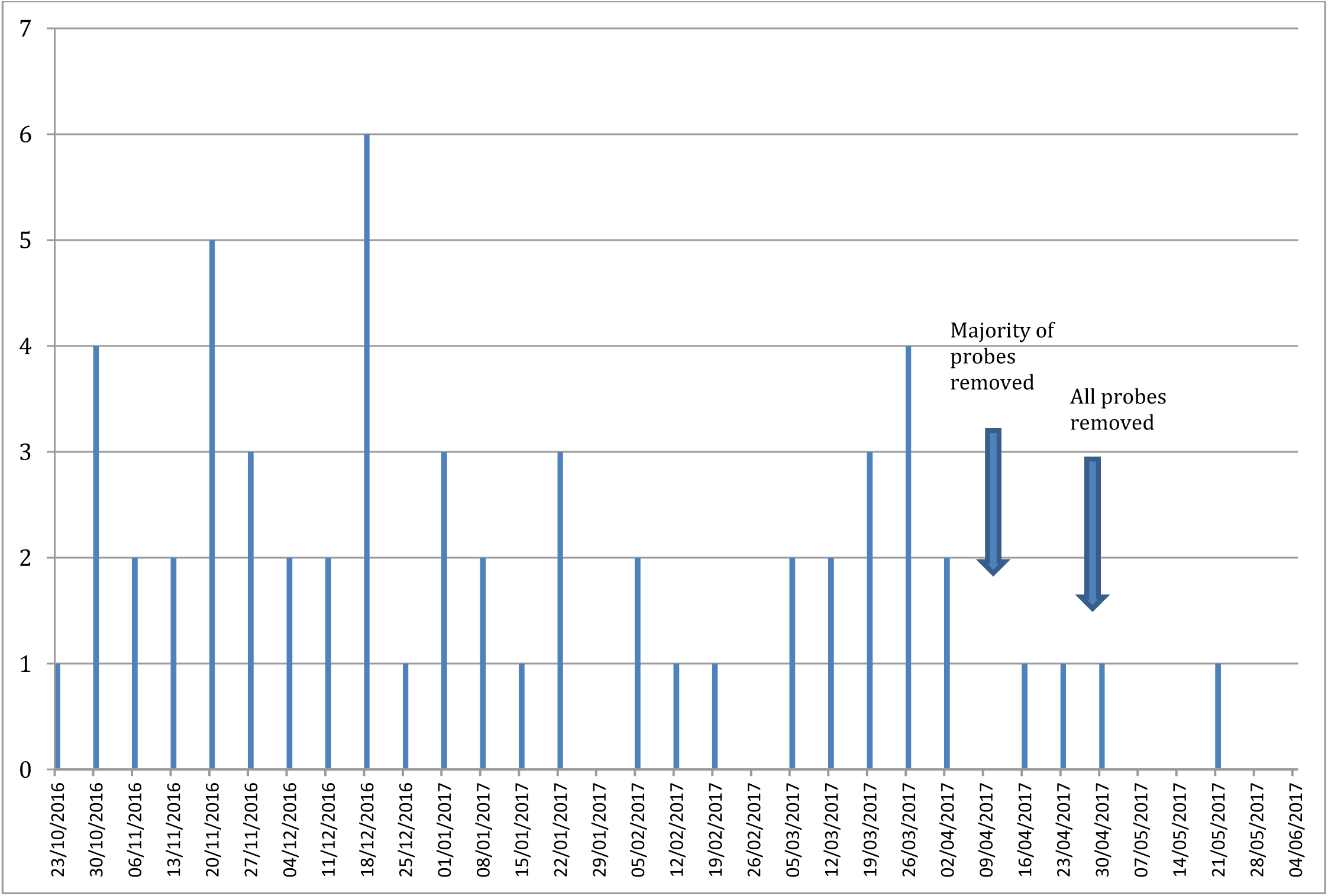
Weekly *Candida auris* new acquisition rate on the NITU. y axis: number of patients with newly identified *C. auris* positive screening samples x axis: Date in weekly increments

### Environmental screening

There were three phases of environmental screening; November 2016, February 2017, and April 2017, with follow-up samples as indicated. A total of 128 samples were obtained. Sampling methods and results are detailed in Table 1. Briefly, *C. auris* detection in the general environment was low (one settle plate positive on one sampling point). However, the organism was detected from multiuse patient monitoring equipment (skin surface temperature probes and a pulse oximeter) and a patient hoist.

**Table 1:** Results of environmental sampling

| Site | Date | Sampling method | Culture (Sabouraud Dextrose Agar +/- Chloramphenicol) | Enrichment (Brain heart) |
| --- | --- | --- | --- | --- |
| Pulse oximeter | 23/11/16 | Swab in liquid transport medium | **1/1 *C. auris*** |  |
| Dynamic mattresses x4 | 23/11/16 | Swab in liquid transport medium | 4/4 No growth |  |
| Curtains | 23/11/16 | Contact plate | No growth |  |
| Blood pressure cuff | 23/11/16 | Swab in liquid transport medium | No growth |  |
| Air x 16§ | 24/11/16 | SabC Settle plates | **1 colony *C. auris*^** |  |
|  |  |  | 2 x 1 colony *C. guilliermondii* |  |
|  |  |  | 1 colony *C. albicans* |  |
|  |  |  | 12/16 plates No growth |  |
| Pulse oximeters x 12 | 29/11/16 | Swab in liquid transport medium | 12/12 No growth |  |
| Central ventilation ceiling tile | 29/11/16 | Swab in liquid transport medium | No growth |  |
| Waste water x 4 sink P traps | 03/02/17 | 50mls from P trap | 4/4 No growth |  |
| Sink surface x 4 | 03/02/17 | Polywipe plus 5ml saline | 4/4 No growth |  |
| Ceiling fan/tile | 03/02/17 | Polywipe plus 5ml saline | 1/1 No growth | 1/1 No growth |
| Computer keyboard* | 03/02/17 | Polywipe plus 5ml saline | 4/4 No growth | 4/4 No growth |
| Suction apparatus* | 03/02/17 | Polywipe plus 5ml saline | 4/4 No growth | 4/4 No growth |
| Bedrails* | 03/02/17 | Polywipe plus 5ml saline | 4/4 No growth | 4/4 No growth |
| Trolley** | 03/02/17 | Polywipe plus 5ml saline | 2/2 No growth | 2/2 No growth |
| Ventilator/controls *** | 03/02/17 | Polywipe plus 5ml saline | 2/2 No growth | 2/2 No growth |
| 40 surfaces**** | 04/04/17 | Contact plate | 38/40 no growth (see below) |  |
| Temperature probe x 1 | 04/04/17 | Contact plate | **1/1 *C. auris*** |  |
| Patient mobile hoist | 04/04/17 | Contact plate | **1/1 *C. auris*** |  |
| Twelve surfaces# | 04/04/17 | Cotton-tipped swabs and malt extract broth | 12/12 no growth | 12/12 No growth (malt extract broth) |
| Air¥ | 04/04/17 | Air Ideal (sampling at 100 L/min) | No growth |  |
| Air¥ | 04/04/17 | Coriolis sampler (sampling at 300 L/min) | No Growth |  |
| Temperature probe x 1^^ | 15/04/17 | Swab in liquid transport medium | No growth |  |
| Temperature probe x 10 | 16/05/17 | Contact plates and enrichment culture | 10/10 No growth | **4/10 *C. auris*** |

§Two settle plates were placed in 4 bedspacesbed spaces for 5 hours during a day shift, and repeated during a nightshift.
^ plate initially in an empty bedspace; a colonised patient was moved into the bed during the period of observation.
¥ Air samplers were positioned in a negative bed space; a positive bed space; a positive side room; an empty (previously positive) bed space and at the doctors/nursing station. Can assume that at the time of sampling, the number of *C. auris* in the air was < 1 cfu/m3 of air sampled.
\*From 4 bedspaces, 2 occupied by colonised patients.
\*\*From 2 bedspaces occupied by colonised patients.
\*\*\*From 2 unoccupied bedspaces.
\*\*\*\* From 2 bedspaces, 1 occupied by a colonised patient

Following the finding of *C. auris* on a skin surface temperature probe using a contact plate (<u>http://www.philips.co.uk/healthcare/product/HC989803162641/skinsurface-temperature-probe-sensor</u>), the probes were withdrawn from use on April 11th. However, during the annual leave of a senior nurse, the probes came back into use, and acquisitions continued (Figure 1). All probes were withdrawn from the unit on April 24^th^, and handed over to the Infection, Prevention and Control team for culture. Ten probes were cultured, of which four probes had been in recent use on the ward, and six were in storage or broken/damaged. *C. auris* was cultured from four probes (on enrichment in brain heart infusion broth). It is not known if these were the four probes in recent use.

## Discussion

Recommended infection, prevention and control (IPC) measures to control outbreaks of *C. auris* include patient contact isolation, and enhanced cleaning with chlorine based products (3,4). Additional measures taken on the NITU were ‘decluttering’ to facilitate cleaning, the reduction of bedside stock of equipment and other items, and removal of fans and forced air convection blankets. Despite the implementation of these intensive IPC measures, the outbreak in the Oxford NITU has been prolonged and with a high acquisition rate, with detection of colonisation or infection in 20% (57/280) patients during the screening period described.

On the NITU, continuous patient temperature monitoring with a re-usable skin surface temperature probe in the axilla was part of routine care for ventilated patients. Cleaning of probes between patients is by means of quaternary ammonium compound pre-prepared wipes (Clinell Universal Sanitising Wipes). The temperature probes are difficult to clean, with a rubber two-layer sheath protecting the distal end of the wire adjacent to the sensor (Figure 2). We provide evidence that *C. auris* can be recovered from ‘cleaned’ skin surface temperature probes supporting the hypothesis that re-usable equipment is a risk factor for colonisation/infection with this organism. Removal of skin surface temperature probes from the unit appears to reduce the acquisition rate (Figure 1).

**Figure 2.**
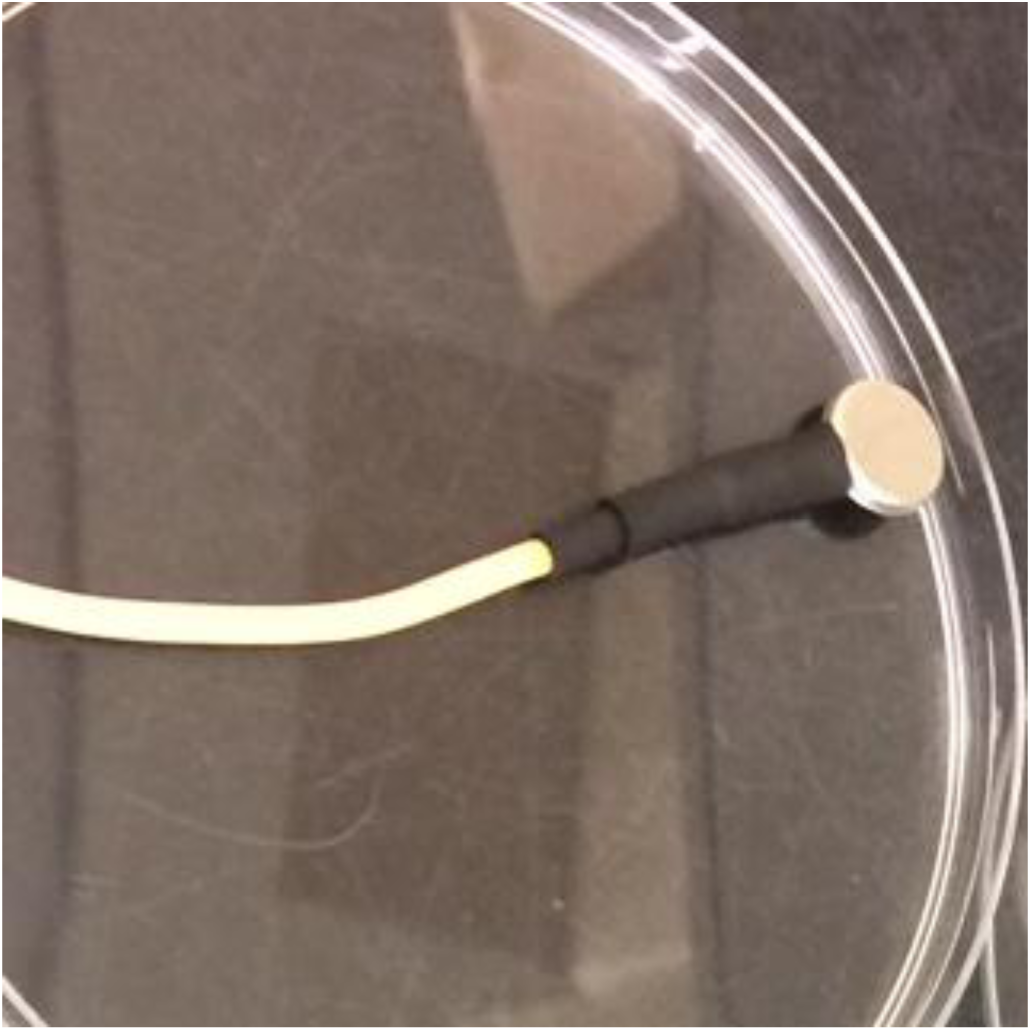
Skin surface temperature probe.

Our initial observations will be further investigated by conducting a cohort study examining risk factors for colonisation and infection using the Infections in Oxford Research database (5). We continue the three-times weekly screening for *C. auris* on all patients in the NITU.

## Acknowledgements

The research was supported by the National Institute for Health Research Health Protection Research Unit (NIHR HPRU) in Healthcare Associated Infections and Antimicrobial Resistance at Oxford University in partnership with Public Health England (PHE). The views expressed are those of the author(s) and not necessarily those of the NHS, the NIHR, the Department of Health or Public Health England.

